# Abnormal X Chromosome Inactivation in Females with Major Psychiatric Disorders

**DOI:** 10.1101/009555

**Authors:** Baohu Ji, Kerin K. Higa, John R. Kelsoe, Xianjin Zhou

**Affiliations:** Department of Psychiatry University of California San Diego 9500 Gilman Drive La Jolla, CA 92093

## Abstract

Bipolar disorder, major depression and schizophrenia are severe brain disorders. No biological hallmark has been identified for any of these disorders. Here, we report that abnormal X chromosome inactivation (XCI) often presents in lymphoblastoid cells of female patients with different major psychiatric disorders in the general population. X chromosome inactivation is well preserved in human lymphoblastoid cells. *XIST*, *KDM5C,* and other X-linked genes are over-expressed in the lymphoblastoid cells of female patients, suggesting an abnormal XCI. Trimethylation of lysine 27 on histone 3 (H3K27me3) is significantly increased at both *XIST* and *KDM5C* gene loci. We found that *XIST* and *KDM5C* expression can be used as a potential diagnostic hallmark for major psychiatric disorders in a large sub-population of female patients. Preliminary studies also suggest an increased *XIST* expression in postmortem brains from female patients with schizophrenia, bipolar disorder, and major depression. An increased gene dosage from some X-linked genes may contribute to the development of psychiatric disorders, as functional disomy of partial X chromosome have been suggested to cause mental retardation and other developmental abnormalities. Additionally, patients with Klinefelter syndrome (XXY) or Triple X syndrome (XXX) frequently display psychiatric disorders due to an extra X chromosome. Mutation of the *KDM5C* gene was reported to cause X-linked syndromic mental retardation. Our studies suggest that abnormal X chromosome inactivation could play a causal role in development of major psychiatric disorders in females. Correction of abnormal X chromosome inactivation may prevent and/or cure major psychiatric disorders in a sub-population of female patients in the future.

## Introduction

Schizophrenia, major depression and bipolar disorder are severe brain disorders. The biological basis of these major psychiatric disorders has been little understood. Our previous studies found that inhibition of protein translation may contribute to pathogenesis of major psychiatric disorders in a rare Scottish family (Ji et al., 2014). Conversely, excessive protein translation has been suggested in fragile X syndrome and autism (Santoro et al., 2012). We speculate that abnormal protein translation may contribute to a wide range of mental disorders not only in rare families, but also in the general population. It is impossible to directly measure protein translation activity in the human brain. However, there are some genes that can affect protein translation in both lymphocytes and neurons; for example, mutation of *FMR1* causes excessive protein translation in both lymphoblastoid cells and the brain (Bhakar et al., 2012; Gross and Bassell, 2012). Therefore, we analyzed protein translation activity in psychiatric patients’ lymphoblastoid cells in hope of validating our hypothesis and finding more susceptibility genes. We observed a significantly larger variation in protein translation activity in the patients’ lymphoblastoid cells than in the cells of healthy controls. Surprisingly, all variations in protein translation activity came from the female patients. These findings prompted us to investigate functions of the X chromosome in the female patients’ lymphoblastoid cells. We examined X chromosome inactivation (XCI) by quantifying expression of *XIST* and several other X-linked genes in the lymphoblastoid cells of patients with different psychiatric disorders. A preliminary study of *XIST* expression was conducted in the postmortem brains of patients with schizophrenia, bipolar disorder, and major depression. Finally, chromatin immunoprecipitation was performed to examine abnormalities of epigenetic modifications of the X chromosome.

## Results

### Abnormal protein translation activity in lymphoblastoid cells from patients with severe mania and psychosis

Protein translation activity was examined in the lymphoblastoid cells of psychiatric patients. To synchronize cell growth, lymphoblastoid cells were first serum-starved for 8 h before re-stimulated with a normal concentration of serum. Protein translation activity was measured 8 h after serum re-stimulation using the SUnSET (surface sensing of translation) method (Ji et al., 2014; Schmidt et al., 2009). Different protein translation activities were observed between cell lines (Figure 1A). After analyzing 26 healthy controls and 28 patients with severe mania and psychosis, we found a significantly larger variation in protein translation activity in the patients’ lymphoblastoid cells (Figure 1B). To rule out whether variations in protein translation activity measured by SUnSET may come from differential puromycin uptake between cell lines, we quantified the intracellular concentrations of puromycin using a modified quantification method (Fujiwara et al., 1982). We selected two cell lines (B25 and B27) with a marked difference in SUnSET (Figure 1C). The concentration of intracellular puromycin in each cell line was quantified by the effectiveness of its cell lysate to block the interactions between anti-puromycin antibodies and immobilized puromycin-containing proteins on the Western blot membrane. Western blot signal was gradually decreased after adding an increasing amount of cell lysate to the blocking solution (Figure 1C). There was no difference between the blocking activities of B25 and B27 cell lysate, indicating that there is a comparable concentration of intracellular puromycin between the two cell lines. We conclude that different SUnSET intensities between B25 and B27 likely come from differential activities of protein synthesis. We also conducted SUnSET experiments in asynchronous proliferating lymphoblastoid cells. There was a high correlation of SUnSET intensities between the proliferating cells and the synchronized cells (data not shown). No correlation was found between individuals’ age and protein translation activity (Figure S1A). When the SUnSET data were sub-grouped according to gender, however, we found that all variation in protein translation activity came from the female patients (Figure 1D). To avoid potential bias in the normalization of SUnSET data between gels, we re-ran females’ SUnSET on the same Western blot gel. Larger variation in protein translation activity was confirmed in the female patients (Figure S1B).

**Figure 1.**
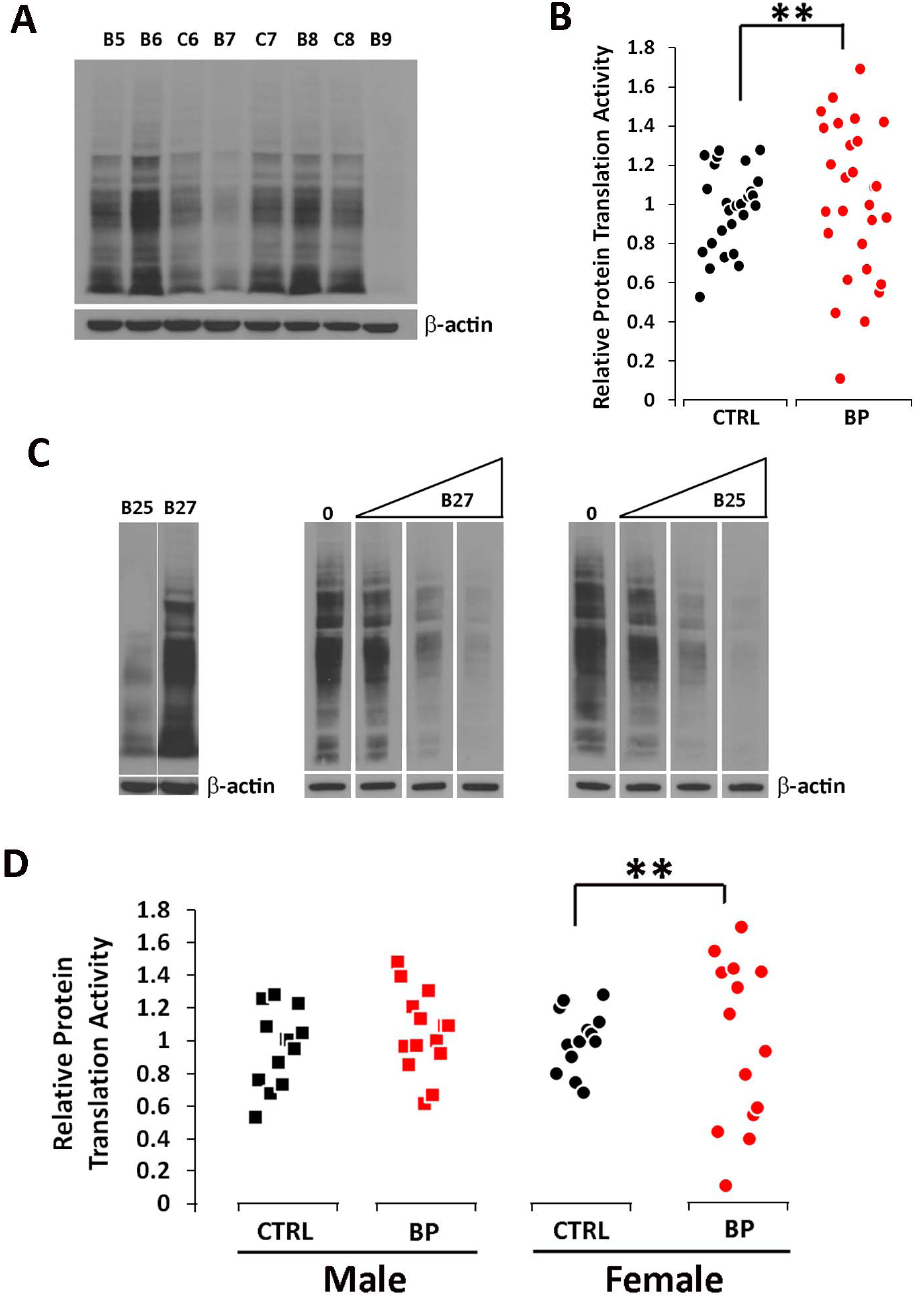
Variation of protein translation activity in patients’ lymphoblastoid cells. **(A)** Protein translation activity was measured in individual lymphoblastoid cell lines using SUnSET. Sample loading was shown by β-actin protein expression after stripping the membrane. C1-C26: healthy European Caucasian controls; B1-B28: European Caucasian bipolar patients with mania and psychosis. **(B)** SUnSET intensity of individual samples was normalized against β-actin. To avoid differential overall intensity between the gels, the intensity of each sample is presented as a percentage of the average intensity of all samples from the same gel. A significantly larger variation of protein translation activity was observed in the patients than in the controls (p < 0.01, *F*-test). Each dot represents a human subject (black=healthy controls; red=patients). CTRL: controls; BP: bipolar patients with mania and psychosis. **(C)** Comparison of intracellular puromycin concentrations. Two cell lines (B25 and B27) with a marked difference in SUnSET activity were selected. Equal amounts of puromycin-labled protein were loaded on each lane. B25 and B27 cell lysates were used to block interaction between membrane-bound puromycin-labeled proteins and anti-puromycin antibodies in the Western blot blocking solution. Increase of either cell lysate had a similar magnitude of blocking effect, suggesting a comparable concentration of intracellular puromycin despite their remarkable difference in SUnSET. **(D)** SUnSET data were analyzed according to gender. There was no difference between male controls and male patients. A significantly larger variation of protein translation activity was observed in female patients (p < 0.01, *F*-test). Each dot represents a human subject (black=healthy controls; red=patients). CTRL: controls; BP: bipolar patients with mania and psychosis.

### XCI dysregulation in female patients with severe mania and psychosis

Restriction of abnormal protein translation activity to the female patients’ lymphoblastoid cells prompted us to investigate potential dysregulation of sex chromosome functions. We examined XCI in female lymphoblastoid cells by quantifying expression of *XIST* (Brown et al., 1992), *TSIX* (Migeon et al., 2002), *FTX* (Chureau et al., 2002), and *JPX* (Chureau et al., 2002) genes. All of these genes produce non-coding RNAs (ncRNA) and are localized at the core of the X chromosome inactivation center (XIC). Their mouse orthologs play central roles in establishing XCI (Brockdorff et al., 1992; Chureau et al., 2011; Lee et al., 1999; Tian et al., 2010). We found that expression of *XIST* is significantly higher (*p* = 0.001, FDR corrected for multiple comparisons) in the female patients than in the female controls (Figure 2A). In contrast to high *XIST* expression, *TSIX* expression is significantly lower (*p* < 0.01, FDR corrected for multiple comparisons) in the patients’ cells (Figure 2B). We also observed a trend of high expression of *FTX* (*p* < 0.1, FDR corrected for multiple comparisons) in the patients (Figure 2C), but no difference was found in *JPX* expression (Figure 2D). In mouse, *Xist* expression is negatively regulated by *Tsix*, but positively regulated by *Ftx* and *Jpx* (Chureau et al., 2011; Lee et al., 1999; Tian et al., 2010). Consistent with the negative role of mouse *Tsix*, human *TSIX* expression is lower in the patients’ cells that display higher *XIST* expression. However, there is only a modest linear negative correlation between human *XIST* and *TSIX* expression (Figure 2E). *TSIX* may play different roles in human from its mouse ortholog *Tsix* (Migeon et al., 2002). There is a modest negative correlation between human *TSIX* and *FTX* expression (Figure S2A), consistent with the opposite roles proposed for mouse *Tsix* and *Ftx* genes in the regulation of *Xist* expression. Taken together, altered expression of *XIST* and its possible regulator genes within XIC indicates an abnormal XCI in the lymphoblastoid cells of the female patients.

**Figure 2.**
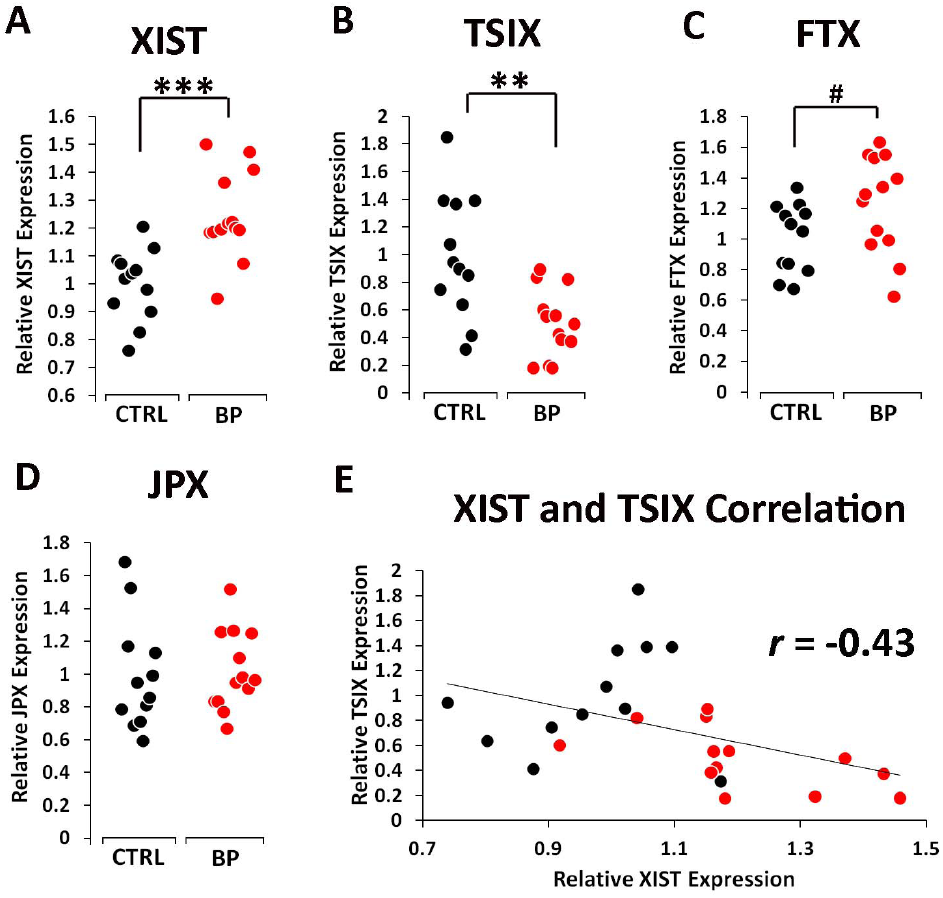
Abnormal expression of *XIST* and its regulator genes in X chromosome inactivation center. Each dot represents a human subject (black=healthy controls; red=patients). CTRL: controls; BP: bipolar patients with mania and psychosis. **(A)** A significantly higher *XIST* RNA expression was observed in the patients than in the controls (t(23)=-4.23, p = 0.001, unpaired, two-tailed student’s *t*-test, FDR corrected for multiple comparisons). **(B)** Consistent with its negative role, there is a significantly lower expression of *TSIX* in the patients (t(23)=3.43, p < 0.01, unpaired, two-tailed student’s *t*-test, FDR corrected for multiple comparisons). **(C)** Consistent with its positive role, a trend of high expression of *FTX* was detected in the patients (t(23)=-2.00, p < 0.1, unpaired, two-tailed student’s *t*-test, FDR corrected for multiple comparisons). **(D)** No difference was observed in *JPX* expression between the controls and patients (t(23)=-0.24, ns, unpaired, two-tailed student’s *t*-test, FDR corrected for multiple comparisons). **(E)** A modest negative linear correlation was present between human *XIST* and *TSIX* expressions in all samples (Pearson’s coefficient, *r* = -0.43).

Abnormal XCI may affect expression of other X-linked genes beyond XIC. We randomly selected a few genes that are either completely inactivated by or escapees from XCI. *PGK1*, *G6PD* and *HPRT1* genes have been known to be inactivated by XCI, and *KDM5C*, *KDM6A* and *RPS4X* genes are the well-established escapees from XCI in human lymphoblastoid cells (Johnston et al., 2008). We found that expression of *KDM5C* is significantly higher (*p* < 0.05, FDR corrected for multiple comparisons) in the female patients than in the female controls (Figure 3A). There is a trend of high expression of *KDM6A*, *PGK1* and *G6PD* genes (p < 0.1, uncorrected) in the patients’ cells before correction of multiple comparisons (Figure 3B, 3C, 3D). No difference was found in expression of either *HPRT1* or *RPS4X* gene between the patients and the controls (Figure S2B, S2C). *KDM5C* and *KDM6A* genes encode different histone H3 lysine-specific demethylases that play key roles in chromatin remodeling (Iwase et al., 2007; Lan et al., 2007; Lee et al., 2007; Tahiliani et al., 2007). A strong positive correlation (Pearson’s coefficient, *r* = 0.88) was observed between *KDM5C* and *KDM6A* expression (Figure 3E). Interestingly, *XIST* and *KDM5C* expression also display a strong positive correlation (Figure 3F)(Pearson’s coefficient, *r* = 0.78). We explored whether a combination of *XIST* and *KDM5C* expression can be used as a potential diagnostic marker to separate the patients from the controls. According to calculation of reference range in clinical blood tests (William J. Marshall, 2008), we calculated the reference ranges of *XIST* and *KDM5C* expression in the control females to define a “normal” distribution. The upper limit of the reference range is defined as 1.96 standard deviations above the mean (dashed line, Figure 3F). Out of 13 patients, 8 have *XIST* and/or *KDM5C* expression above their reference ranges.

**Figure 3.**
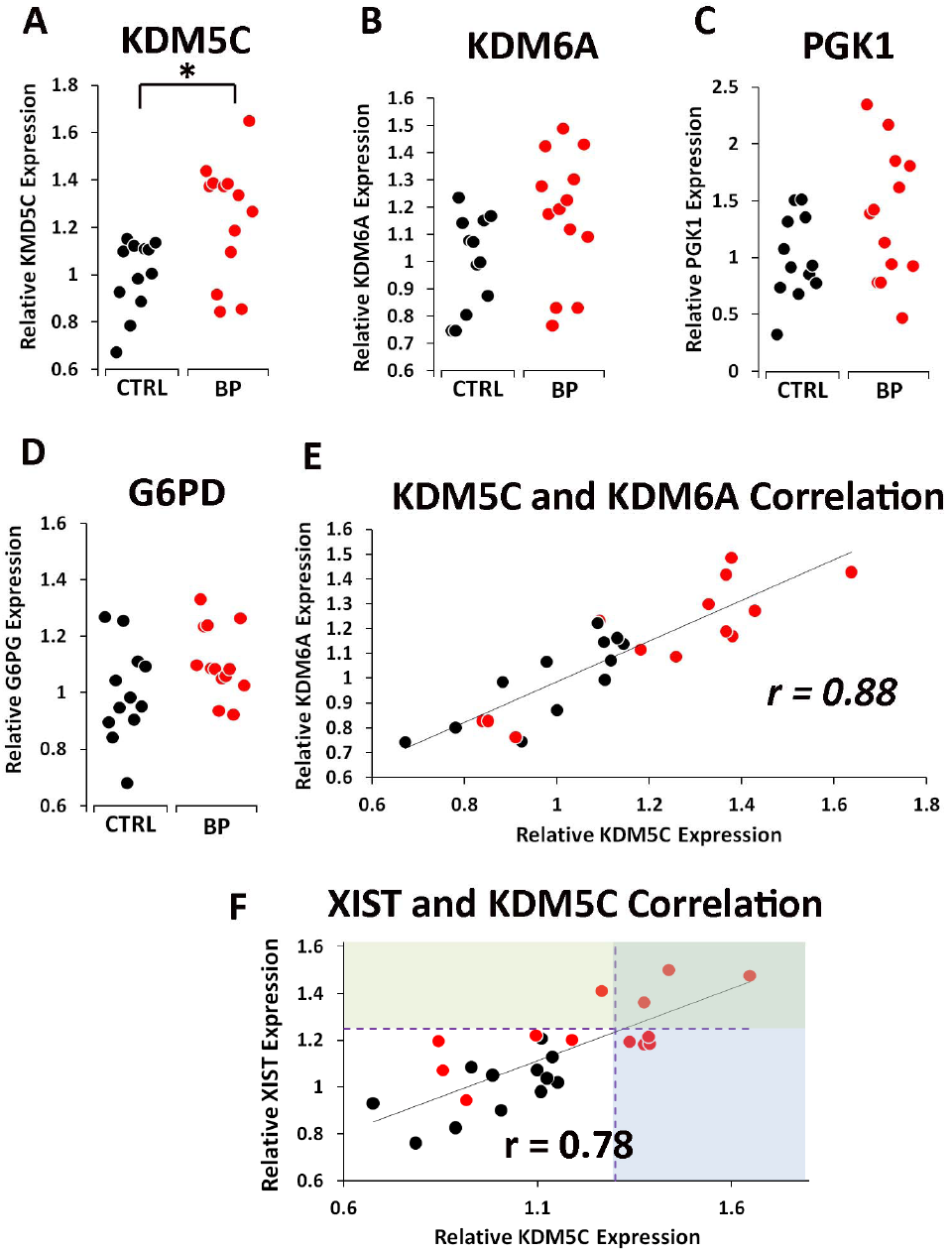
Increased expression of genes outside of X chromosome inactivation center. Each dot represents a human subject (black=healthy controls; red=patients). CTRL: controls; BP: bipolar patients with mania and psychosis. **(A)** A significantly higher *KDM5C* expression (t(23)=-2.89, p < 0.05, unpaired, two-tailed student’s *t*-test, FDR corrected for multiple comparisons) was observed in the patients. There was a trend of high expression in *KDM6A* **(B)**(t(23)=-2.00, p < 0.1, unpaired, two-tailed student’s *t*-test), *PGK1* **(C)**(t(23)=-1.84, p < 0.1, unpaired, two-tailed student’s *t*-test), *G6PD* **(D)**(t(23)=- 1.89, p < 0.1, unpaired, two-tailed student’s *t*-test) in the patients before correction of multiple comparisons. **(E)** A high correlation between *KDM5C* and *KDM6A* RNA expressions was observed in all samples (Pearson’s coefficient, *r* = 0.88). **(F)** There is a high correlation between *KDM5C* and *XIST* RNA expressions (Pearson’s coefficient, *r* = 0.78). Reference ranges of *XIST* and *KDM5C* expression in healthy controls were calculated as an interval between which 95% of values of a reference group fall into. The upper limit of reference range is defined as 1.96 standard deviations above the mean. *XIST* and *KDM5C* expression above the upper limit of their reference ranges are shaded with light green and light purple respectively.

To investigate whether female patients’ medication histories may potentially alter expression of *XIST*, *TSIX, KDM5C* and *KDM6A* genes in their later established lymphoblastoid cells, we examined expression of these genes in lymphoblastoid cells after treatment of the antipsychotics clozapine (CLZ) and the mood stabilizer valproic acid (VPA). None of these genes displayed altered expression after a combined treatment of CLZ and VPA (Figure S3A, S3B, S3C, S3D). These data suggest that female patients’ medication histories (if any) are unlikely to have any effect on the expression of *XIST*, *KDM5C* and other genes in their lymphoblastoid cells. Additionally, expression of *KDM5C* and *KDM6A* genes was not altered in the lymphoblastoid cells of male patients in comparison with the healthy male controls (Figure S4), supporting that patients’ medication history has no effect on expression of X-linked genes in their lymphoblastoid cells. Neither *XIST* nor *KDM5C* expression is affected by age (Figure S5A, S5B). We investigated whether abnormal XCI may be responsible for more variation in protein translation activity in the female patients’ lymphoblastoid cells. There is no correlation between *XIST* expression and protein translation activity (SUnSET) (Figure S5C). A weak correlation between *KDM5C* expression and SUnSET was observed (Figure S5D). We speculate that other modifier genes on autosomes, which conceivably vary between individuals, may also be involved in the regulation of protein translation.

### XCI dysregulation in female patients with recurrent major depression

We are particularly interested in the finding of abnormal XCI in the lymphoblastoid cells of some bipolar female patients with mania and psychosis. To investigate whether abnormal XCI presents in a different psychiatric disorder, we examined expression of *XIST*, *TSIX*, *KDM5C, KDM6A, PGK1,* and *G6PD* genes in the lymphoblastoid cells of female patients with recurrent major depression. These genes were chosen because they displayed significant or trend of increases of expression in patients with mania and psychosis. We found that expression of *XIST* is significantly higher (*p* < 0.001, FDR corrected for multiple comparisons) in the female patients with recurrent major depression than in the female healthy controls (Figure 4A). In contrast to significant reduction of *TSIX* expression in the female patients with mania and psychosis, down-regulation of *TSIX* expression was not statistically significant in the lymphoblastoid cells of female patients with recurrent major depression (Figure 4B). It is possible that the sample size of the patients may be too small to have a sufficient statistical power to detect a smaller difference. Expression of *KDM5C* is also significantly higher (*p* < 0.001, FDR corrected for multiple comparisons) in the patients with recurrent major depression (Figure 4C). Consistent with a trend of high *KDM6A* expression in female patients with mania and psychosis, we observed a significantly higher expression of *KDM6A* (p < 0.05, FDR corrected for multiple comparisons) in the lymphoblastoid cells of female patients with recurrent major depression (Figure 4D). However, no difference was found in either *PGK1* or *G6PD* expression between the female patients and healthy controls (Figure S6A, S6B). As expected, a strong positive correlation (Pearson’s coefficient, *r* = 0.85) was observed between *KDM5C* and *KDM6A* expression (Figure 4E). *XIST* expression is also highly correlated with *KDM5C* expression (Pearson’s coefficient, *r* = 0.85)(Figure 4F). After calculating the reference ranges of *XIST* and *KDM5C* expression in the healthy controls, 8 out of 10 patients have *XIST* expression above the upper limit of its reference range.

**Figure 4.**
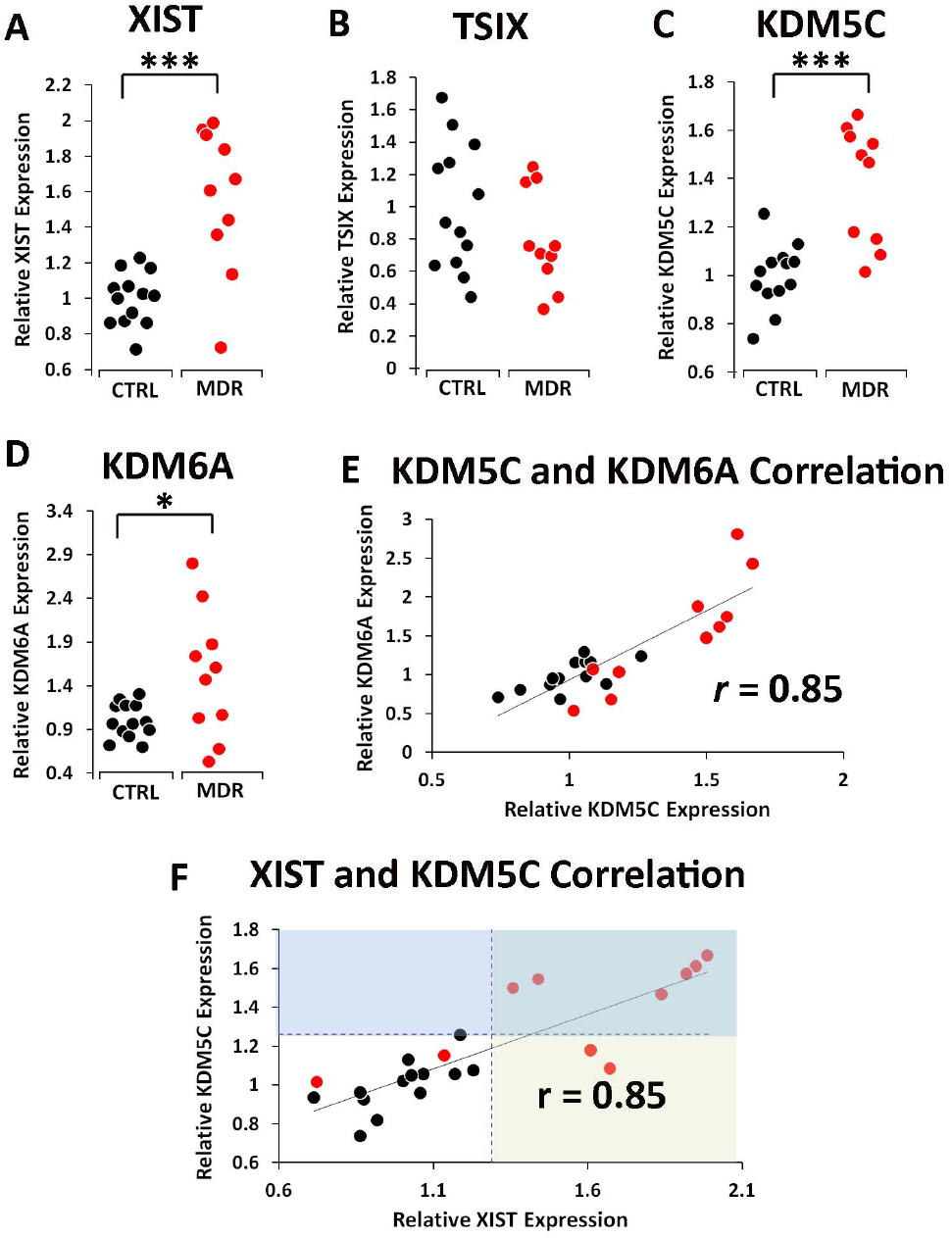
Abnormal XCI in patients with recurrent major depression. Each dot represents a human subject (black=healthy controls; red=patients). CTRL: controls; MDR: patients with recurrent major depression. **(A)** There is a significantly higher *XIST* expression (t(21)=-4.62, p < 0.001, unpaired, two-tailed student’s *t*-test, FDR corrected for multiple comparisons) in the patients than in the controls. **(B)** Down-regulation of *TSIX* expression was not statistically significant (t(21)=1.35, ns, unpaired, two-tailed student’s *t*-test, FDR corrected for multiple comparisons). A significantly higher expression of *KDM5C* **(C)**(t(21)=-4.79, p < 0.001, unpaired, two-tailed student’s *t*-test, FDR corrected for multiple comparisons) and *KDM6A* **(D)**(t(21)=-2.51, p < 0.05, unpaired, two-tailed student’s *t*-test, FDR corrected for multiple comparisons) were observed. **(E)** Consistent with previous results, a high correlation between *KDM5C* and *KDM6A* RNA expression was found (Pearson’s coefficient, *r* = 0.85). **(F)** As expected, there is a high correlation between *KDM5C* and *XIST* RNA expressions (Pearson’s coefficient, *r* = 0.85). Reference ranges of *XIST* and *KDM5C* expression in healthy controls were calculated as previously described. *XIST* and *KDM5C* expression above the upper limit of their reference ranges are shaded with light green and light purple respectively.

We next examined expression of KDM5C and KDM6A proteins in female patients with recurrent major depression. Western blot analyses confirmed a significantly higher level of KDM5C protein (p < 0.01) in the patients’ cells than in the controls’ cells (Figure 5A, 5B). A modest correlation between the levels of *KDM5C* RNA and KDM5C protein was observed (Pearson’s coefficient, *r* = 0.53)(Figure 5C). Conceivably, protein expression is not solely controlled by the level of RNA expression (Schwanhausser et al., 2011). Western blot detected a weaker signal for KDM6A protein (Figure S6C). A significantly higher level of KDM6A proteins was also observed in the patients (p < 0.05)(Figure S6D). No correlation between *KDM6A* RNA and KDM6A protein expression was detected (Figure S6E). We also measured protein translation activity in the female patients’ cells using SUnSET. A significantly larger variation of protein translation activity was also observed in the lymphoblastoid cells of female patients with recurrent major depression (data not shown).

**Figure 5.**
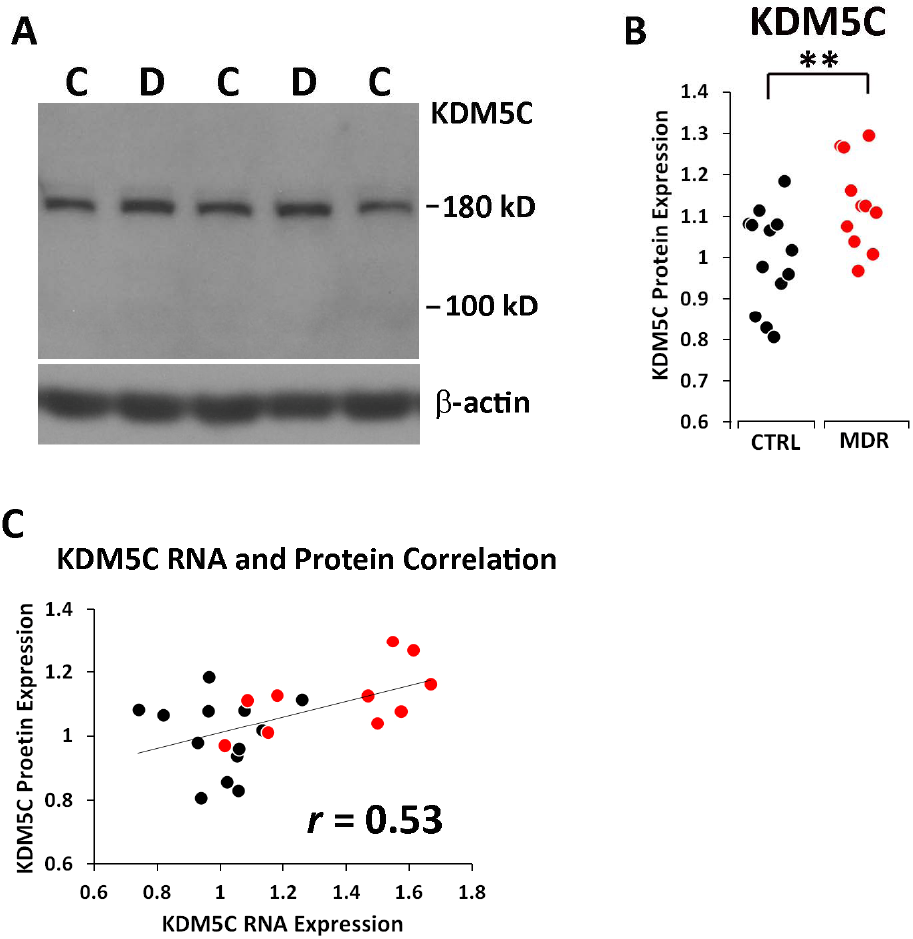
Increased expression of KDM5C protein in patients’ lymphoblastoid cells. Each dot represents a human subject (black=healthy controls; red=patients). CTRL: controls; MDR: patients with recurrent major depression. During the experiments, we found an additional patient cell line from -80 oC freezer, which was included in the Western blot analyses. **(A)** Western blot analyses of KDM5C protein expression in the lymphoblastoid cells of female patients with recurrent major depression. A single band at 180 kD, the calculated size of human KDM5C protein, was detected. C: controls; D: recurrent major depression. β-actin was used as an internal control for normalization. **(B)** Consistent with increased RNA expression, a significantly higher KDM5C protein expression was found in the patients (t(22)=-, p < 0.01, unpaired, two-tailed student’s *t*-test). **(C)** A modest correlation between *KDM5C* RNA and KDM5C protein expression was observed across all samples (Pearson’s coefficient, *r* = 0.53).

### A steady-state level of *XIST* expression

To investigate whether cell passages may alter expression of *XIST*, *KDM5C,* and *KDM6A* genes, we collected batches of cells from different cell passages and analyzed expression of these genes. The level of *XIST* expression is very stable between different cell passages (Figure 6A). There is a high correlation of *XIST* expression (Pearson’s coefficient, *r* = 0.88) in individual cell lines between the two different cell passages. Despite more variation in *KDM5C* expression (Figure 6B), there is still a strong correlation between the two different cell passages (Pearson’s coefficient, *r* = 0.6). In contrast to *XIST* and *KDM5C* expression, *KDM6A* expression appeared to be more affected by variations from different cell cultures (Figure S7A).

**Figure 6.**
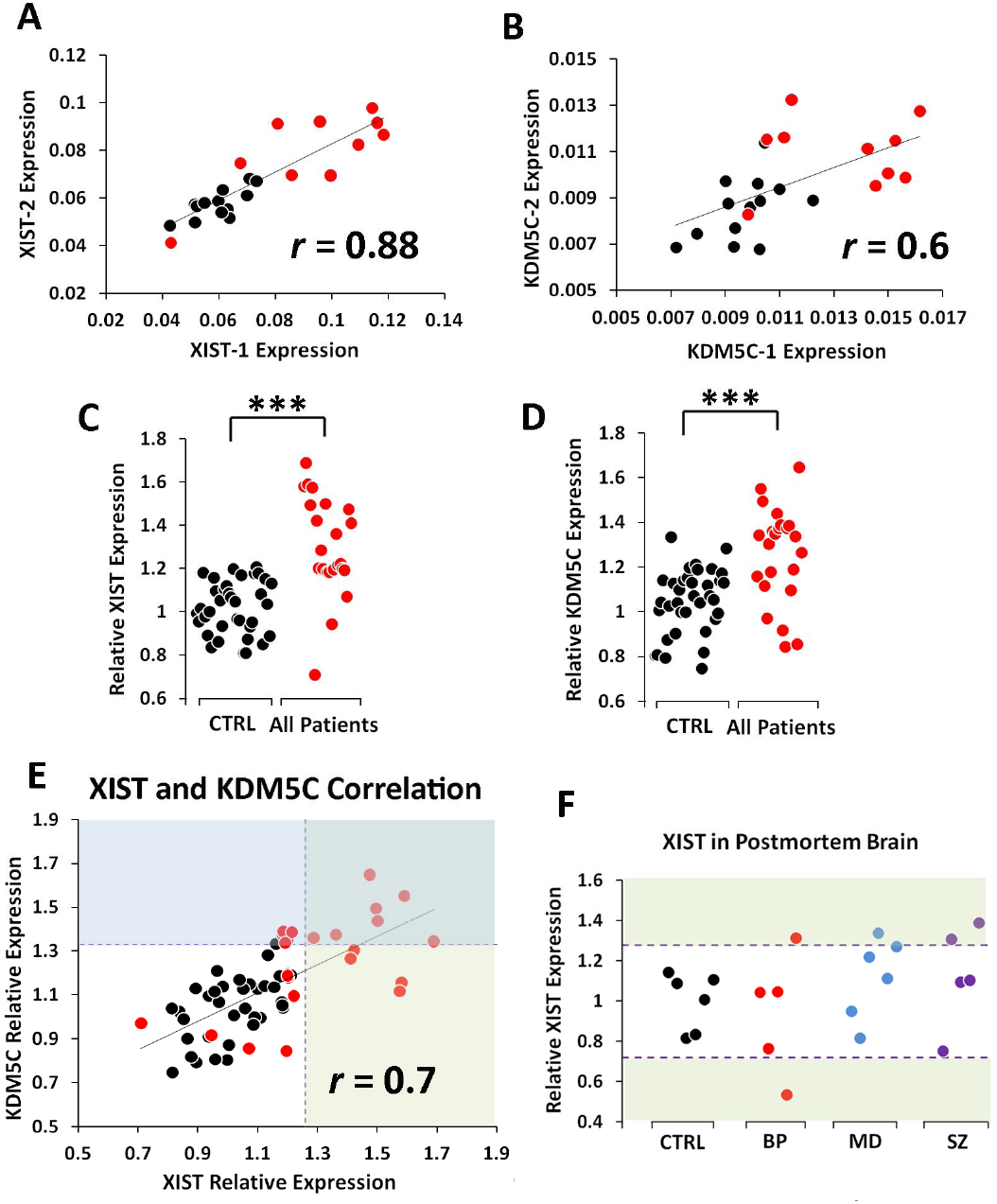
A steady-state level of *XIST* expression between different cell passages and across different ethnic backgrounds. **(A)** *XIST* and *KDM5C* gene expression was examined in the first batch of lymphoblastoid cells (*XIST*-1 and *KDM5C*-1). After more than a month of continuous cell culture, the same set of genes was examined again in the second batch of cells with different cell passages (*XIST*-2 and *KDM5C*-2). *XIST* and *KDM5C* expression was only normalized against β-actin without being further presented as a percentage relative to the mean of the controls in order to compare absolute amount of RNA transcripts between different cell passages. A high correlation of *XIST* expression was observed between the two batches of cells (Pearson’s coefficient, *r* = 0.88). **(B)** There is a relatively strong correlation in *KDM5C* expression between the two batches (Pearson’s coefficient, *r* = 0.6). **(C)** To enlarge the sample size of healthy female controls, 23 more healthy females with mixed ethnic background were included in analysis of *XIST* expression. Since both depression and mania patients display the same abnormal XCI, they were combined in analyses. Significantly higher *XIST* expression (t(57)=-6.06, p < 10^-6^, unpaired, two-tailed student’s *t*-test, FDR corrected for multiple comparisons) was found in the combined group of patients than in the combined group of the healthy controls. **(D)** *KDM5C* also displayed significantly higher expression (t(57)=-4.51, p < 10^-4^, unpaired, two-tailed student’s *t*-test, FDR corrected for multiple comparisons) in the combined group of patients. **(E)** As expected, a high correlation between *XIST* and *KDM5C* expression was observed (Pearson’s coefficient, *r* = 0.7). Reference ranges of *XIST* and *KDM5C* expression in the combined healthy controls were calculated as previously described. *XIST* and *KDM5C* expression above the upper limit of their reference ranges are shaded with light green and light purple respectively. **(F)** *XIST* expression was quantified in postmortem human brains. Several patients have high *XIST* expression above the upper limit of its reference range calculated from the healthy control brains. The upper limit (1.279) of the reference range of *XIST* expression in brain is very close to that (1.26) of the reference range calculated from the combined controls’ lymphoblastoid cells.

To address potential lack of representativeness of the small sample size of the female controls, we enlarged the control sample size by including more healthy females. Because we ran out of female European Caucasian controls, we analyzed expression of *XIST*, *KDM5C* and *KDM6A* genes in a group of healthy females with mixed ethnic backgrounds. There is no difference in either *XIST* or *KDM5C* expression between the European Caucasian female controls and the female controls with mixed ethnic backgrounds (Figure S7B, S7C). A slightly higher expression of *KDM6A* gene was observed in the controls with mixed ethnic backgrounds (Figure S7D). Considering that *KDM6A* expression is more susceptible to variations of different cell cultures, it is not surprising that its expression can be affected by different ethnic backgrounds. Since there is no ethnic effect on either *XIST* or *KDM5C* expression, we combined the two groups of healthy female controls together. We have demonstrated that female patients with either mania and psychosis or recurrent major depression display over-expression of both *XIST* and *KDM5C* genes. Therefore, we combined the two groups of female patients together and examined their XCI again. As expected, *XIST* expression displays a narrow distribution in the combined female controls (Figure 6C). A significantly higher *XIST* expression was observed in the combined female patients (*p* < 10^-6^, FDR corrected for multiple comparisons) than in the combined female controls. Females with Triple X syndrome (XXX), although rare, have a high *XIST* expression. We did not find any of our controls or patients with triple X chromosomes (Figure S7E). There is also a significantly higher *KDM5C* expression in the combined female patients (*p* < 10^-4^, FDR corrected for multiple comparisons)(Figure 6D). With an increased sample size of healthy controls, more accurate reference ranges were calculated for both *XIST* and *KDM5C* expressions. 16 out of 23 patients displayed high *XIST* (11 patients) and/or *KDM5C* (12 patients) expression above their reference ranges (Figure 6E). Considering that the female patients and the female controls were randomly collected from the general population, it is likely that a significantly large sub-population of female patients from the general population may have abnormal XCI.

In a limited number of postmortem human brains, we examined expression of *XIST*, *TSIX, KDM5C*, and *KDM6A* genes in the cortical RNA of female patients with schizophrenia, bipolar disorder, and major depression (kindly provided by Stanley Medical Research Institute). Due to small sample sizes, we did not expect to detect group differences in gene expression. We rather focused on identifying individuals who may display abnormal XCI. Despite the small sample size of female controls’ postmortem brains, the distribution range of *XIST* expressions is very similar to the one calculated from the combined controls’ lymphoblastoid cells. In the postmortem brains, the upper limit (1.279) of the reference range of *XIST* expression is close to the one (1.26) calculated from the combined controls’ lymphoblastoid cells, confirming little variation of *XIST* expression among healthy individuals. Several female patients display higher *XIST* expression above the upper limit (1.279) of its reference range in the postmortem brains (Figure 6F). We noticed that one bipolar patient displayed a lower *XIST* expression. *TSIX* expression in the 6 patients who have the highest *XIST* expression is significantly lower than that in the rest of population (Figure S7F), supporting potentially abnormal XCI in these patients’ brains. In contrast to strong correlations between different gene expressions in lymphoblastoid cells, no correlation between *XIST*, *KDM5C*, and *KDM6A* expressions was observed (data not shown). Many confounding factors, such as tissue and cell heterogeneity, RNA degradation, disease conditions etc, could complicate RNA expression analysis in the postmortem brains. More human postmortem brains are needed to confirm the findings in future.

### Increased trimethylation of lysine 27 on histone 3 (H3K27me3) on the X chromosome in female patients’ lymphoblastoid cells

H3K27me3 is particularly enriched on the inactive X chromosome because of *Xist*-dependent recruitment of Polycomb repressor complex 2 (PRC2) that methylates lysine 27 on histone 3 (Maenner et al., 2010; Plath et al., 2003; Zhao et al., 2008). Over-expression of *XIST* may increase H3K27me3 on the inactive X chromosome in female patients’ lymphoblastoid cells. However, we observed over-expression of *XIST*, *KDM5C* and other X-linked genes in the patients. H3K27me3 was therefore examined around the promoters of *XIST* and *KDM5C* genes using chromatin immunoprecipitation. Out of 6 randomly selected sites at the promoter of *XIST* gene, *XIST* (1) and *XIST* (3) sites have significantly more abundant H3K27me3 in the patients’ lymphoblastoid cells than in the controls’ lymphoblastoid cells (Figure 7A). A trend of more H3K27me3 was observed at *XIST* (4) in the patients. Consistent with more H3K27me3 on the *XIST* gene, *KDM5C* gene also displayed more abundant H3K27me3 at *KDM5C* (1) in the patients’ lymphoblastoid cells (Figure 7B). H3K27me3 enrichment at some loci appears to correlate well with the level of gene expression (Figure 7C, 7D, 7E). These data provided further support to the abnormal XCI in female patients’ lymphoblastoid cells.

**Figure 7.**
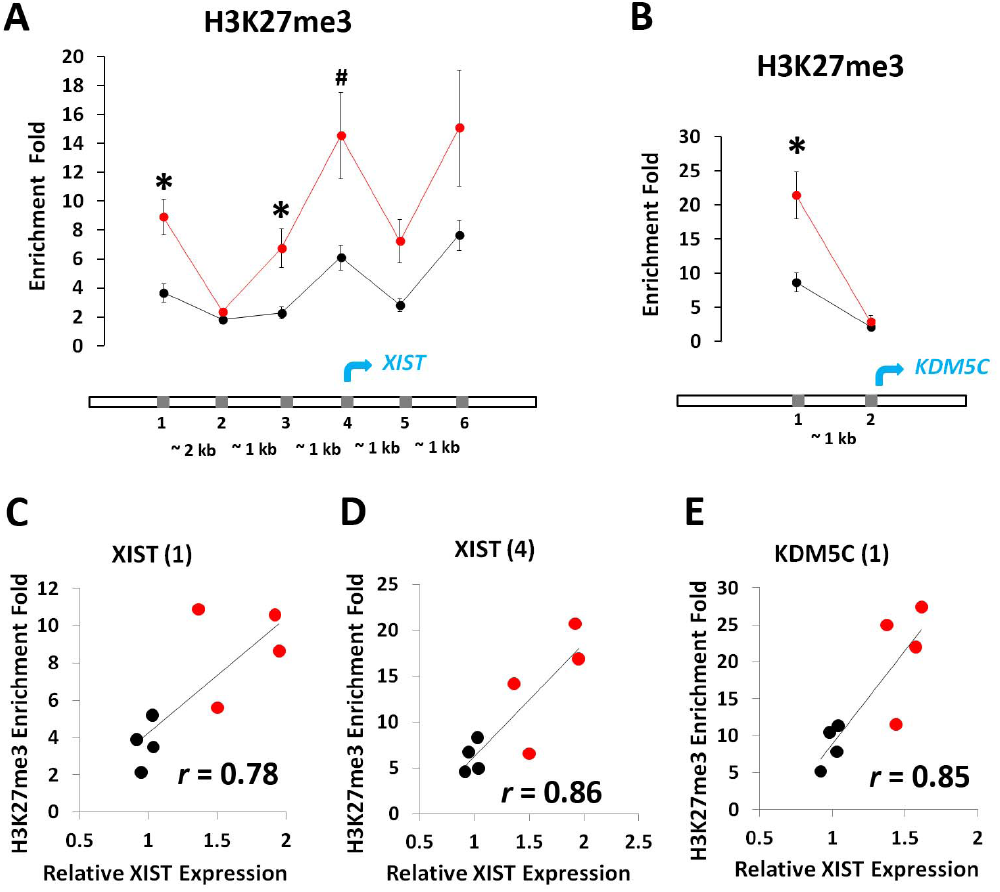
Enrichment of H3K27me3 at the promoters of *XIST* and *KDM5C* genes. Lymphoblastoid cell lines from 4 female controls (black) and 4 female patients (red) with either recurrent major depression or mania and psychosis were selected for chromatin immunoprecipitation experiments. Six sites, separated by ∼1 kb (except a 2 kb between 1 and 2) around the promoter of *XIST* gene, were examined by ChIP-QPCR. Two sites around the promoter of *KDM5C* gene were also examined. **(A)** Significantly more H3K27me3 was observed in the patients’ lymphoblastoid cells at *XIST* (1) (t(6)=-3.83, p < 0.05, unpaired, two-tailed student’s *t*-test, FDR corrected for multiple comparisons) and *XIST* (3) (t(6)=-3.2, p < 0.05, unpaired, two-tailed student’s *t*-test, FDR corrected for multiple comparisons). A trend of more H3K27me3 at *XIST* (4) (t(6)=-2.7, p < 0.1, unpaired, two-tailed student’s *t*-test, FDR corrected for multiple comparisons) was also observed. **(B)** H3K27me3 is significantly more enriched at *KDM5C* (1) (t(6)=-3.39, p < 0.05, unpaired, two-tailed student’s *t*-test, FDR corrected for multiple comparisons) in female patients’ lymphoblastoid cells. High correlations between enrichment of H3K27me3 and *XIST* expression were detected at *XIST* (1) **(C)** and *XIST* (4) **(D)**. A modest correlation was also observed at *XIST* (3). **(E)** In *KDM5C* gene, there was a high correlation between enrichment of H3K27me3 and *KDM5C* expression at *KDM5C* (1). Error bar: SEM. (* p < 0.05, # p < 0.1)

### Discussion

Lymphoblastoid cells have been used to identify differential gene expressions as potential biomarkers for psychiatric disorders. However, these studies have been questioned for their relevance to psychiatric disorders due to lack of justification of commonality between lymphoblastoid cells and the brain. XCI is unique. It occurs early during embryogenesis in all somatic cells to ensure the same gene dosage from X chromosome between males and females (Lee and Bartolomei, 2013). XCI is very stable and well preserved in human lymphoblastoid cells (Johnston et al., 2008; Zhang et al., 2013), providing a convenient cell model to study XCI effects in human disease (Amir et al., 2000). Therefore, we consider abnormal XCI observed in patients’ lymphoblastoid cells to reflect the same abnormality of XCI in all somatic cells including the brain. Our preliminary studies of a limited number of postmortem human brains support the conclusion.

In mouse, *Xist* is a master gene in the initiation of XCI (Kay et al., 1993; Penny et al., 1996; Plath et al., 2002). *Tsix*, *Ftx*, and *Jpx* genes, localized in X chromosome inactivation center (XIC), have been demonstrated to regulate expression of *Xist* (Chureau et al., 2011; Lee et al., 1999; Tian et al., 2010). In human lymphoblastoid cells, we observed a significantly higher *XIST* expression in the patients with either mania and psychosis or recurrent major depression. Alteration of *TSIX* and *FTX* expression was less robust, but consistent with the regulatory roles proposed for their mouse orthologs. However, there may be species difference in the mechanisms of regulation; for example, human *TSIX* was reported to be co-expressed with *XIST* from the same inactivated X chromosome in contrast to mouse *Tsix* that expresses only in the active X chromosome (Migeon et al., 2002). Alteration of gene expression caused by abnormal XCI spreads beyond XIC in the female patients. Out of 6 randomly selected genes outside of XIC, *KDM5C* and *KDM6A* genes displayed significantly higher expression in female patients’ lymphoblastoid cells. Disinhibition of X-linked gene expression suggested a deficient XCI in the female patients’ lymphoblastoid cells. Unexpectedly, we observed more abundant H3K27me3, an epigenetic silencing marker on the inactive X chromosome (Plath et al., 2003), at both *XIST* and *KDM5C* genes in the female patients’ lymphoblastoid cells. It is possible that H3K27me3 level may also be altered at other X-linked genes. How can we reconcile excessive H3K27me3 at these genes with their over-expression in the patients’ lymphoblastoid cells? One possible scenario could be that over-expression of *XIST* may result from disinhibition of X-linked genes by deficient XCI. Excessive *XIST* RNA recruits more PRC2 to increase H3K27me3 at *XIST*, *KDM5C,* and other genes on the inactive X chromosome. Although H3K27me3 is a negative cue that can be recognized by other repressors such as PRC1 to silence gene expression, H3K27me3 does not alter chromatin structure per se to suppress gene expression (Margueron and Reinberg, 2011). Increased H3K27me3, therefore, may not necessarily be sufficient to suppress expression of X-linked genes in female patients’ lymphoblastoid cells. Although H3K27me3 is in general a repressive marker, it may play an active role in expression of some set of genes (Young et al., 2011). There is another possibility that over-expression of *XIST* may cause, rather than a consequence of, deficient XCI. For example, excessive H3K27me3 generated by *XIST* overexpression may interfere with establishment of insulators (Phillips-Cremins and Corces, 2013; Van Bortle et al., 2012; Weth et al., 2014) and thereby compromise the organization of inactive X chromosome structure to disinhibit expression of X-linked genes. In addition to *XIST*-PRC2 in the regulation of H3K27me3, KDM6A that demethylates H3K27me3 was increased in the patients’ lymphoblastoid cells. With a strong correlation with *KDM6A* expression, *KDM5C* that demethylates H3K4me3 was also over-expressed in the female patients’ lymphoblastoid cells. Dysregulation of histone 3 epigenetic modifications appears complex in the females with abnormal X chromosome inactivation. The X chromosome-wide RNA-seq and ChIP-seq analyses may offer better mechanistic understanding of the abnormal X chromosome inactivation in the patients’ lymphoblastoid cells in the future. Taken together, increased *XIST* expression could be either a consequence or a causal factor to induce abnormal XCI. It is possible that *XIST* overexpression may also be a compensatory response to deficient XCI in female patients’ lymphoblastoid cells.

We observed a trend of higher expression of *PGK1* and *G6PD* genes in bipolar patients with mania and psychosis, consistent with over-expression of *KDM5C* and *KDM6A*. Considering more than 1,000 genes on the X chromosome, more X-linked genes may increase their expression in patients’ lymphoblastoid cells. Could increased gene dosage from the X chromosome contribute to development of psychiatric disorders? It is well known that gene over-dosage causes human diseases, e.g. Down syndrome. For the X chromosome, there are reports of functional disomy of partial X chromosome in the pathogenesis of mental retardation and other developmental abnormalities (Lahn et al., 1994; Sanlaville et al., 2005). Patients with Klinefelter syndrome (XXY) or Triple X syndrome (XXX) frequently display a variety of psychiatric disorders that are presumably caused by over-dosage of the X-linked escapee genes (DeLisi et al., 1994; DeLisi et al., 2005; Otter et al., 2010). We did not find any Triple X syndrome patients in our samples. In Triple X syndrome, all escapee genes on the extra inactive X chromosome are presumably over-expressed. In the female patients (XX) we analyzed, however, not all escapee genes (e.g. *RPS4X*) were over-expressed. Increased expression of X-inactivated genes was also observed in some patients with mania and psychosis. Therefore, there may be differences in the over-dosage of X-linked genes between Triple X syndrome and the female patients (XX) with abnormal X chromosome inactivation. Therefore, we posit that over-dosage of some X-linked genes could cause major psychiatric disorders in female patients with a normal karyotype (46, XX). Interestingly, mutation of *KDM5C* gene was reported to cause X-linked syndromic mental retardation (Tahiliani et al., 2007); however, functions of its over-expression remain unknown.

It is interesting that *XIST* has a steady-state level of high expression and correlates well with *KDM5C* gene expressions in individual cell lines across different cell passages. These observations suggest that human *XIST* may still have a functional role beyond the establishment of XCI, presumably in maintenance of XCI. To support our hypothesis, mouse *Xist* was demonstrated to be essential for maintenance of XCI (Yildirim et al., 2013). However, the role of *XIST/Xist* expression is less understood in the maintenance than in the initiation of XCI. It is unlikely that *TSIX, FTX,* and *JPX* genes play a major role in the maintenance of *XIST* expression because of lack of robust correlations between expression of *XIST* and expression of any of these genes.

Our studies suggest that there is an abnormal XCI in female patients’ lymphoblastoid cells, which may disinhibit expression of X-linked genes. We do not assume that the same set of X-linked genes affected by abnormal XCI will display the same alteration of expression between the patients’ lymphoblastoid cells and brains since cell-specific regulators will also contribute to alteration of their expression. Therefore, consequence of abnormal XCI on gene expression in patients’ brain remains unknown, which is likely more relevant to development of major psychiatric disorders in females.

Abnormal XCI has been observed in patients with either mania and psychosis or recurrent major depression. How could the abnormality of XCI result in markedly different clinical symptoms? One explanation could be that psychiatric disorders are not distinct individual diseases, but a spectrum of mental disorders. For example, some patients display both mania and depression. The same chromosome translocation associates with schizophrenia, major depression, and other psychiatric disorders in a Scottish family (St Clair et al., 1990). Another explanation is that abnormal XCI may interact with other modifier genes on autosomes and environment to result in different clinical symptoms. In psychiatric genetics, no single genetic locus has been found to carry a high risk and affect a significantly large portion of the general population of patients. It has been assumed that men and women share common genetic risk factors. The X chromosome is often omitted in GWAS studies (Wise et al., 2013). Surprisingly, we showed that a large subpopulation of female psychiatric patients from the general population may actually have a different genetic cause or pathway (XCI) from men. We would argue that this subpopulation of female patients may be better defined biologically as a “XIST disorder” rather than being categorized into different psychiatric disorders according to their clinical symptoms. Identification of a subpopulation of “XIST disorder” with a common genetic and biological causality could accelerate scientific research, and provide objectivity in clinical diagnosis as well as a biological target for development of treatment.

Many questions remain to be answered. We found abnormal protein translation in the lymphoblastoid cells of female patients. Does abnormal XCI contribute to abnormal protein translation activity in female patients’ lymphoblastoid cells? We did not observe a correlation between *XIST* expression and protein translation activity. If factors other than XCI are responsible for protein translation abnormalities, why do male patients’ lymphoblastoid cells not display abnormal protein translation activity? We observed a weak correlation between *KDM5C* expression and protein translation activity, indicating that abnormal XCI may indirectly contribute to dysregulation of protein translation activity in female patients’ cells. Because regulation of protein translation activity is very complex and sensitive to culture conditions in patients’ lymphoblastoid cells, it is more difficult to use its measurement as a potential biomarker. Therefore, we shifted our focus to the investigation of abnormal XCI that could be a biological cause as well as a potential biological hallmark for major psychiatric disorders in some females.

The number of genetic factors regulating *XIST/Xist* expression in initiation of XCI, a fundamental biological process during embryogenesis, remain incompletely understood (Lee, 2011). It was reported that mouse *Xist* expression quickly reaches a high level to initiate XCI and maintains the same high level across later development (Buzin et al., 1994) and between different adult tissues (Kay et al., 1993; Ma and Strauss, 2005). Our cell culture studies of human *XIST* across different cell passages also support a steady-state level of *XIST/Xist* expression. If the level of *XIST* expression is a relative stable character of an individual, measurement of *XIST* expression in lymphoblastoid cells or any other somatic cells conceivably may provide an objective test to diagnose “XIST disorder”, and even at childhood to predict the risk of developing mental illness at adolescence or later. To fully evaluate the feasibility of *XIST* as a potential diagnostic marker, however, studies on more controls and patients are needed. It will also be interesting to know whether abnormal XCI presents in females with neurodevelopmental disorders (e.g. autism) or other human diseases. If abnormal XCI is a major genetic and biological cause, correction of abnormal XCI may prevent and/or cure major psychiatric disorders in this subpopulation of female patients in future.

## Materials and Methods

### Lymphoblastoid cell culture

Studies on human lymphoblastoid cells were conducted under a University of California San Diego IRB-approved protocol. All lymphoblastoid cell lines were kindly provided by Dr. John R. Kelsoe at UCSD. The cells were in their early passages (< 20 passages)(Oh et al., 2013). Demographic information of the subjects is summarized in Table SI. All female patients except one have a family history with one or more relatives suffering from psychiatric disorders. There is no significant difference in age between healthy controls and patients when blood was taken to generate lymphoblastoid cells by transformation with Epstein-Barr virus (EBV). A second group of healthy female controls with mixed ethnic backgrounds were included to expand the size of controls and to examine potential effect of ethnic background on *XIST* expression. Lymphoblastoid cell lines were cultured in Roswell Park Memorial Institute (RPMI) 1640 medium containing 10% fetal bovine serum (Life Technologies, CA) with penicillin and streptomycin (Life Technologies, CA) at 37°C in a humidified atmosphere of 5% CO2.

### Protein translation activity

Lymphoblastoid cells were serum-starved for 8 h to synchronize cell growth. 8 h after addition of serum, SUnSET was conducted to measure instant protein translation activity as described (Ji et al., 2014). Patient and control samples were loaded alternately on PAGE gels regardless of age and gender. Because of differential overall intensities between different gels in western blot analyses, the signal of each sample was first normalized against the average intensity of all samples from the same gel. After normalization, samples were analyzed according to diagnosis, gender, and age. To confirm a larger variation of protein translation activity in female patients, female samples were re-run on the same Western blot gel to rule out potential bias introduced by normalization between gels.

### Quantification of intracellular puromycin concentration

Puromycin-labeled protein used to quantify intracellular concentrations of puromycin in lymphoblastoid cells was prepared as follows: HEK293T cells were labeled with puromycin for 20 minutes (Ji et al., 2014). Cells were collected and sonicated in passive lysis buffer (Promega, WI) with 1X protease inhibitor cocktail (P8340, Sigma). Protein concentration was measured with Bradford (Abs 595nm) method with Coomassie Plus Protein Assay (Thermo Scientific, IL). B25 and B27 lymphoblastoid cells were incubated with puromycin for 20 min. The cells were then spun down to remove supernatant and washed twice before lysis. Quantification of intracellular puromycin concentration by Western blot: 15 ug puromycin-labeled proteins were loading on 4-20% gradient Tris-Glycine gel. After electrophoresis, proteins were transferred onto PVDF membranes. After 1 hour blocking with 5% of nonfat milk in TBST buffer (pH 7.5, 10 mM Tris-HCl, 150 mM NaCl, and 0.1% Tween 20) at room temperature, the membrane were cut into individual lanes. Each lane was incubated overnight with anti-puromycin antibody (dilution at 1:200,000; 12D10, Millipore) in the presence of either B25 or B27 cell lysate. After washing three times, the blot was incubated with horseradish peroxidase (HRP)-conjugated anti-mouse IgG (dilution at 1:10,000, Cell signaling, MA) for 1.5 hr at room temperature. Pierce ECL Western blot substrate (Pierce, IL) was used to visualize the signal. Quantification of Western blot bands was conducted using Image J.

### RNA from postmortem human brains

We received 57 BA7 cortical RNA samples out of 60 brains from Stanley Medical Research Institute (SMRI), consisting of 15 each diagnosed with schizophrenia, bipolar disorder, or major depression, and 15 unaffected controls. 21 out of 57 samples were females. The four groups were matched by age, sex, race, postmortem interval, pH, hemisphere, and mRNA quality (http://www.stanleyresearch.org/dnn/Default.aspx?tabid=196).

### Quantitative real-time RT-PCR

Total RNA was extracted from lymphoblastoid cells with TRIzol reagent (Invitrogen, Carlsbad, CA). Primers were selected to amplify exons without alternative splicing in order to quantify all RNA isoforms. All PCR primers were first confirmed to specifically amplify the target cDNA genes without background before being used for QPCR. cDNA was synthesized from 5ug of total RNA using Superscript III First-Strand Synthesis System (Invitrogen, Carlsbad, CA) with random hexamers. SYBR Green real-time PCR was used to quantify relative expression of all genes with a comparative Ct method. The standard curve of PCR amplification was examined. All amplifications had R^2^ >0.99. Amplification efficiency for each pair of primers was determined using known molecular concentration of template DNA. Each sample had 4 amplification replica wells. After amplification, Ct was set in the exponential amplification phase of the curve as recommended by the manufacturer’s protocol (Bio-Rad CFX384). Variation of cycles between amplification replica wells was smaller than 0.5 cycles. β-actin was used as a reference control for normalization. In most cases, relative expression of each gene in each sample was presented as a percentage relative to the mean of the control group. The following primers were used in real-time QPCR:

#### 1. β-actin

Forward primer: 5’ TTCTACAATGAGCTGCGTGTG3’

Reverse primer: 5’ GGGGTGTTGAAGGTCTCAAA3’

#### 2. XIST

Forward primer: 5’ GGATGTCAAAAGATCGGCCC3’

Reverse primer: 5’ GTCCTCAGGTCTCACATGCT3’

#### 3. TSIX

Forward primer: 5’ TGTGCCAGGTAATAGAAACACA3’

Reverse primer: 5’ ACCTGCATTTGTGGGATTGT3’

#### 4. KDM5C

Forward primer: 5’ GGGTCCGACGATTTCCTACC3’

Reverse primer: 5’ GCGATGGGCCTGATTTTCG3’

#### 5. KDM6A

Forward primer: 5’ CGCTTTCGGTGATGAGGAAA3’

Reverse primer: 5’ GTCCTGGCGCCATCTTCAT3’

#### 6. FTX

Forward primer: 5’ GTGATCTGGACAAAGGAACAGG3’

Reverse primer: 5’ CATTTCCCGTGAACACCCG3’

#### 7. G6PD

Forward primer: 5’ CATGGTGCTGAGATTTGCCA3’

Reverse primer: 5’ CCTGCACCTCTGAGATGCAT3’

#### 8. JPX

Forward primer: 5’ ACTGACACTGGTGCTTTCCT3’

Reverse primer: 5’ GGACTCCCAGGCATTGCTAT3’

#### 9. HPRT1

Forward primer: 5’ AGTGATGATGAACCAGGTTATGA3’

Reverse primer: 5’ GCTACAATGTGATGGCCTCC3’

#### 10. PGK1

Forward primer: 5’ GAATCACCGACCTCTCTCCC3’

Reverse primer: 5’ GGGACAGCAGCCTTAATCCT3’

#### 11. RPS4X

Forward primer: 5’ AGGTCCTCTTTCCTTGCCTA3’

Reverse primer: 5’ GAGCAAACACACCGGTCAAT3’

### Western Blot

Lymphoblastoid cells were collected by centrifugation and sonicated in passive lysis buffer (Promega, WI) with 1X protease inhibitor cocktail (P8340, Sigma). Protein concentration was measured using the Bradford (Abs 595nm) method with Coomassie Plus Protein Assay (Thermo Scientific, IL). 50 µg proteins were loaded for Western blot analysis (Ji et al., 2014). Rabbit anti-KDM5C (cat. 39230, dilution at 1:7,500, Active Motif, Carlsbad), rabbit anti-KDM6A (cat. 61516, dilution at 1:7,500, Active Motif, Carlsbad), and mouse anti-β-actin (sc-47778, dilution at 1:10,000, Santa Cruz) were purchased as primary antibodies. Horseradish peroxidase (HRP)-conjugated anti-mouse IgG (dilution at 1:10,000, Cell signaling, MA) or HRP-conjugated anti-rabbit IgG (dilution at 1:10,000, Santa Cruz, CA) were purchased as secondary antibodies. Pierce ECL Western blot substrate (Pierce, IL) was used for chemiluminescence visualization. Quantification of Western blot bands was conducted using Image J.

### Chromatin immunoprecipitation (ChIP)

Lymphoblastoid cell lines from 4 female controls and 4 female patients with high *XIST* expression were selected for examination of H3K27me3 on the inactive X chromosome. 1.5 X 107 cells were harvested from each cell line. Cells were fixed, and chromatin was sheared. Chromatin immunoprecipitation was performed using ChIP-IT High Sensitivity Kit (Active Motif) according to its manual. Mouse monoclonal antibody against H3K27me3 (cat. 61017) was purchased from Active Motif. 6 pairs of primers were designed to amplify immunoprecipitated genomic DNA (each about 150 bp) around the transcription start site of *XIST* gene. 2 pairs of primers were selected to amplify immunoprecipitated genomic DNA around the transcription start site of *KDM5C* gene. Amplification of immunoprecipitated β–actin genomic DNA was used as a reference control to quantify enrichment of H3K27me3 at the promoters of *XIST* and *KDM5C* genes. All primers were first tested for specific amplification of their target DNA without non-specific background before being used for ChIP-QPCR. SYBR Green real-time PCR was used to quantify the amount of DNA immunoprecipitated by anti-H3K27me3 with a comparative Ct method. The following primers were used in ChIP-QPCR:

XIST (1)

Forward primer: 5’ TGGAACACTAGTCAGCCTTAAA 3’

Reverse primer: 5’ CTTTCTGTGCCTGGCTTAGC 3’

XIST (2)

Forward primer: 5’ TGGCACTCTGGAACCTCTTT 3’

Reverse primer: 5’ GGACATGTTCAGAGCTGTGG 3’

XIST (3)

Forward primer: 5’ CTGGAAACTGAAGAGCAAGCT 3’

Reverse primer: 5’ CAGAAGGGAGACAATGGCAAC 3’

XIST (4)

Forward primer: 5’ ACGCCTCTTATGCTCTCTCC 3’

Reverse primer: 5’ ACCCCAAGTGCAGAGAGATC 3’

XIST (5)

Forward primer: 5’ GGGAGAAAAGGTGGGATGGA 3’

Reverse primer: 5’ GCCATGCTAATTCACCCAGG 3’

XIST (6)

Forward primer: 5’ GTCCTAGTTCCTCAGTCCCG 3’

Reverse primer: 5’ AGCAATGCCAAGGGTAAACG 3’

KDM5C (1)

Forward primer: 5’ GCCAAGTACACAACAATTCTGC 3’

Reverse primer: 5’ GGGCTCAAAAGAGTCATGCC 3’

KDM5C (2)

Forward primer: 5’ GGCCCATACCGTTAAGACCA 3’

Reverse primer: 5’ GCACGTTTAAAGGCCAGTGA 3’

β-actin

Forward primer: 5’ TGGTTCTCTCTTCTGCCGTT 3’

Reverse primer: 5’ GCTTTACACCAGCCTCATGG 3’

### Data analysis

All data were first tested for normal distribution using the Kolmogorov–Smirnov test. *F*-tests were used to analyze difference between the standard deviations of the two samples. Unpaired, two-tailed student’s *t*-test was used for statistical analyses between two samples. Correction of multiple comparisons were conducted by using false discovery rate (FDR)(http://www.sdmproject.com/utilities/?show=FDR).

## Acknowledgments

We thank Tanya Shekhtman and Brian Chin for assistance with cell culture, and the Stanley Medical Research Institute for providing RNA samples from postmortem human brains.

## Contributions

X.Z. conceived of, analyzed, and wrote the studies. B.J. and K.H. performed experiments and analyzed data. J.R.K. provided all lymphoblastoid cell lines for the studies.

